# A highly conserved A-to-I RNA editing event within the glutamate-gated chloride channel GluClα is necessary for olfactory-based behaviors in *Drosophila*

**DOI:** 10.1101/2023.05.29.542744

**Authors:** Hila Zak, Eyal Rosenfeld, Tricia Deng, David Gorelick, Shai Israel, Mali Levi, Yoav Paas, Jin Billy Li, Moshe Parnas, Galit Shohat-Ophir

## Abstract

A-to-I RNA editing is an important cellular process that modifies genomically encoded information during transcription, to generate various RNA isoforms from a single DNA sequence. It involves the conversion of specific adenosines in the RNA sequence to inosines by ADAR proteins, resulting in their recognition as guanosines by cellular machinery, and as such plays a vital role in neuronal and immune functions. Given the widespread occurrence of A-to-I RNA editing events across the animal kingdom, with thousands to millions of editing sites found in the transcriptomes of organisms such as flies and humans, identifying the critical sites and understanding their *in-vivo* functions remains a challenging task. Here we show for the first time the physiological importance of a single editing site, found within the extracellular domain of the glutamate-gated chloride channel (*GluClα*), and bridge the gap between its evolutionary conservation across *Drosophila* species and its function in shaping the behavior of adult flies. We used genomic editing to ablate editing at this specific site, such that the endogenous channel harbors only the unedited version and used a battery of behavioral paradigms to analyze the effects on various features of adult behavior. We provide evidence that *GluClα^unedited^* flies exhibit reduced olfactory responses to both appetitive and aversive odors, as well as impaired pheromone-dependent social interactions, and that editing of this site is required for proper processing of olfactory information in olfactory projection neurons. Our findings demonstrate that evolutionary conservation is a useful criterion to pinpoint which of the many editing events has the potential of having a function and pave the path for dissecting the link between RNA modification, neuronal physiology, and behavior.

## Introduction

Adenosine-to-inosine (A-to-I) RNA editing, catalyzed by ADAR enzymes, is a ubiquitous mechanism that generates transcriptomic and proteomic diversity in metazoans^1, 2^. Most of the RNA editing events in mammals occur in non-coding parts of the transcriptome by ADAR1 and serve to prevent aberrant immune responses towards self-dsRNAs^3–6^. Only a small fraction of editing events occur in protein-coding sequences mainly by ADAR2, where the deamination of adenosine to inosines introduces non-synonymous substitutions (known as recoding events) that produce a variety of new protein isoforms from a single DNA sequence^7–12^. The binding of ADAR to double-stranded RNA structures containing target sequence and an editing complementary sequence (ECS) promotes the conversion of specific adenosines within the target sequence into inosines which are subsequently recognized by the translation machinery as guanosine (G)^11^. The majority of recoding events occur within genes that function in neurons, accounting for neuronal deficits exhibited by ADAR2 K.O mice^13^. A central example is an essential recoding event of Glutamine to Arginine within the M2 domain of AMPA receptor that is necessary for the proper function of the channel, the absence of which causes lethality^13–15^. Another functionally important recoding event is found within the calmodulin-binding IQ-domain of the voltage-activated calcium channel (Cav1.3) and functions to regulate the calcium-dependent inactivation kinetics of the channel^16^, that in turn shapes hippocampal plasticity and memory in mice^16, 17^.

Recent advances in RNA sequencing and computational analysis facilitated the identification of millions of editing sites in humans^1, 3, 18–20^, tens of thousands in mice^1^, and thousands in *Drosophila*^9, 21–23^. While the function of RNA editing in *Drosophila melanogaster* resembles that of mammals^8, 24–32^, the majority of editing events in *Drosophila melanogaster* are predicted to cause nonsynonymous protein-coding changes, making it hard to pinpoint which to further study their *in-vivo* function. As a consequence, research in the field was limited to studying the function of few editing events at the biochemical level^12, 33–36^, leaving their physiological function at the whole organism level practically unknown. To bridge this gap, we chose to study a recoding site that shows high evolutionary conservation across several *Drosophila* species and is found within the extracellular domain of glutamate-gated chloride channel (GluClα)^37^. The high evolutionary conservation of the particular recoding of Isoleucine at position 27 to Valine is suggestive of its functional importance^37^.

GluClα is an inhibitory channel in invertebrates that belongs to the Cys-loop ligand-gated family of ion channels^38^, consisting of 5 homologous subunits that are arranged in a radial manner (Fig. 1A). Each subunit has an N-terminal extracellular hydrophilic domain containing a ligand binding pocket, 4 transmembrane domains, and an intracellular segment^39^. Glutamate binding triggers a rapid influx of chloride ions, leading to hyperpolarization of the target cell^38, 40^. *Drosophila* GluClα is broadly expressed in the nervous system^41, 42^, and was shown in recent studies to control homeostatic modulation of presynaptic release at the neuromuscular junction^43^, to regulate visual responses^44, 45^ and the processing of olfactory information^40^. Here we show that recoding of Ile^27^ to Val is necessary for proper olfactory responses, since flies expressing endogenous GluClα in which recoding of Ile^27^ is prevented, exhibit reduced responses to appetitive and aversive odors and to pheromone-based social and sexual responses. The behavioral phenotypes of GluClα unedited flies are associated with altered activity of olfactory projection neurons specifically in the VA1v glomeruli and can be rescued by the expression of a fully edited form of GluClα in projection neurons. Our results demonstrate for the first time the physiological relevance of evolutionary conserved editing events in regulating complex behavior in *Drosophila* and may contribute to understanding the biophysical regulation of Cys-loop receptor-channels.

**Figure 1.**
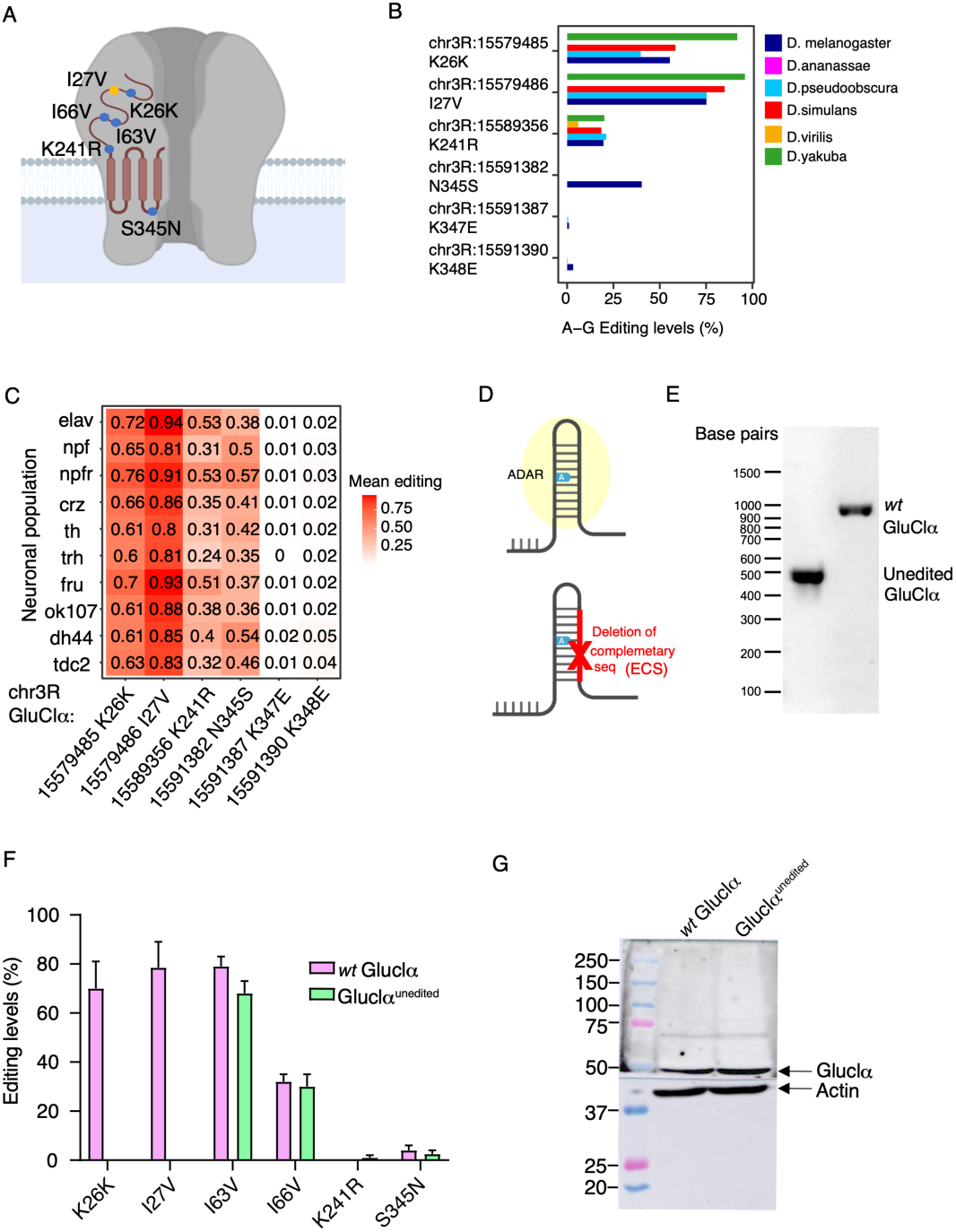
The recoding of Ile^27^→Val is conserved across *Drosophila* species. (A) Schematic representation of the *Drosophila* GluClα receptor and the relative location of its A-to-I editing sites. (B) Relative editing levels of GluClα-editing sites in RNA extracted from various *Drosophila* species. n=2/strain (C) Average editing levels of GluClα sites across various neuronal populations in *Drosophila melanogaster* brain (n=3 for each neuronal population). (D) Schematic illustration of the approach used to generate GluClα unedited flies. ADAR binds to a dsRNA structure formed by base pairing between the exon sequence where editing takes place and a complementary non-coding intronic sequence called the ECS. Deletion of the ECS required for the editing of Ile27 was achieved via CRISPR-Cas 9. (E) PCR validation of ECS deletion in GluClα unedited and *wt* flies. The size of the segment in wild-type flies is ∼900bp whereas in GluClα unedited flies it is ∼490bp (F) Relative editing levels of GluClα sites in GluClα unedited and *wt Drosophila melanogaster (n=* 3 samples/neuronal population). (G) Relative GluClα protein levels in head of GluClα unedited and *wt* flies. Blotting was done with anti-GluClα polyclonal antibodies, and α-Actin as a loading control.

## Results

### The recoding of Ile27 into Val in GluClα is evolutionarily conserved and exhibits high levels of editing

A systematic comparison of conserved RNA editing events across six different *Drosophila* species identified a subset of highly conserved editing sites, suggestive of their functional importance^46^. One of the conserved sites is found in the inhibitory glutamate-gated chloride channel (*GluClα*). The transcript of *GluClα* has six A-to-I editing sites, five of which are classified as nonsynonymous (Figure 1A). Comparing the editing pattern of *GluClα* among the six *Drosophila* species indicated that the recoding event of Ile^27^ located within the extracellular domain of the channel is conserved in *Drosophila melanogaster*, *yakuba*, *simulans,* and *pseudoobscura* with over 75% of the transcripts in all four species are edited (Fig. 1B, Supp. table 1). Next, we analyzed the spatial distribution of the six editing events in *GluClα* across the fly brain using a dataset that we previously generated, containing transcriptomes of nine different neuronal populations including the relative editing levels of thousands of editing sites^47^. We discovered that Ile^27^**→**Val is the most highly edited site in all tested neuronal populations, ranging from 80-94% editing (Fig. 1C, Supp. tables 2 A,B).

The three-dimensional (3-D) structure of the *Drosophila* GluClα receptor is not known. Yet, important structural information can be derived based on homology with other Cys-loop receptors. The editing I**→**V site is situated at position 27 of the immature protein. To locate the editing site in respect to the signal peptide we have analyzed the full-length sequence of the *Drosophila* GluClα subunit using SignalP 6.0^48^. SignalP 6.0 predicted, with very high probability, a signal peptide that includes amino acids 1-22 (Supporting Fig. 1A). As such, the edited I27 is probably the fifth amino acid of the mature protein. We then submitted the full-length amino acid sequence of the immature protein to a secondary-structure prediction by PSIPRED 4.0^49^ and transmembrane (TM) topology prediction^50^ in the PSIPRED server^51^. These predictions support the location of I27 C-terminally (downstream) to the signal peptide and suggest that I27 is located N-terminally to a putative α helix of the *Drosophila* GluClα protein (Supporting Fig. 1B). Very similar secondary-structure prediction obtained with JPred4^52^ corroborated the prediction by PSIPRED 4.0 (not shown). Multiple sequence alignment between the *Drosophila* GluClα subunit and various subunits of anionic Cys-loop receptors (performed by Clustal Omega^53^) indicates that the predicted α helix is aligned with the first α helix (termed α1) of Cys-loop receptor subunits whose 3-D structure was determined at high resolution (Supporting Fig. 1C). In addition it reveals that the editing site (I27**→**V) is indeed located very close to the N-terminus of the mature protein, N-terminally to the first α helix (Supporting Fig. 1C). However, this position is not included in the 3-D structures of the various Cys-loop receptor subunits used for the multiple alignment (Supporting Fig. 1C). Interestingly, in all known 3-D structures of eukaryotic Cys-loop receptors, the N-terminal α helix (α1) proceeds by a loop that connects to the first bstrand (β1). This loop contains a highly conserved amino-acid cluster (Supporting Fig. 1C).

The high ratio of *GluClα* transcripts that undergo editing at this particular site and its evolutionary conservation prompted us to further examine its functional relevance. To this end, we utilized the CRISPR/cas9 system to impair editing at this site by removing the specific editing complementary sequence (ECS, intronic region between positions 15578744-15579171) required for the binding of ADAR (Fig. 1D, E). Comparing the editing pattern of all six sites between *GluClα^unedited^* and matched genetic control showed that the deletion of this particular ECS abolished editing of Ile^27^**→**Val and that of a close-by synonymous site that share the same ECS (Lys^26^**→**Lys), while leaving the editing levels of the other sites intact (Fig. 1F, Supp. table 3). Importantly, the removal of the ECS did not affect protein expression, as can be seen by comparable levels of GluClα protein in protein extracts from *GluClα^unedited^* and control flies (Fig. 1G).

### Recoding of Ile^27^ to Val is necessary for proper olfactory responses

Given the established role of *GluClα* in processing olfactory information^40^, we tested the behavioral responses of *GluClα^unedited^* flies to appetitive and aversive odors. Flies detect odors using olfactory receptor neurons (ORNs) that are in the antennae and maxillary palps. In general, each ORN expresses a single odorant receptor gene^54–57^. ORNs expressing the same receptor send their axons to the same glomerulus in the antennal lobe (AL)^58–60^. In addition, the AL also contains second-order projection neurons (PNs), which receive input from a single glomerulus^40, 61^ and are responsible for delivering odor information to higher brain regions^40, 61^. The AL also contains local neurons (LNs). The majority of LNs are inhibitory GABAergic neurons but about one-third of LNs are inhibitory glutamatergic neurons^40^ which inhibit via *GluClα* both PNs and LNs^40^ (Fig. 2A).

**Figure 2.**
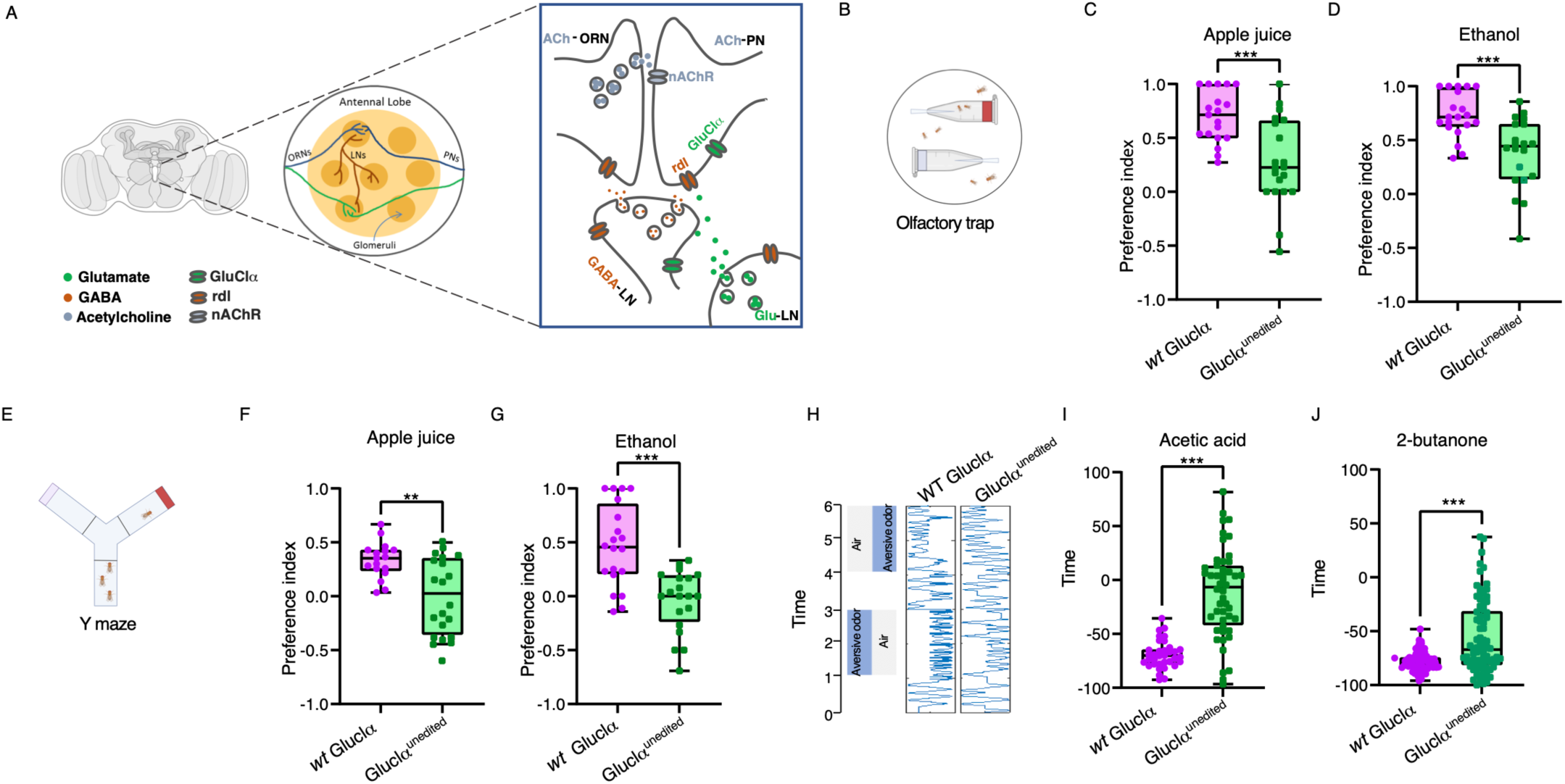
Ile^27^ recoding is necessary for proper olfactory responses. (A) Schematic illustration of GluClα function within the antennal lobe (AL). Projection neurons (PNs) receive input from olfactory receptor neurons (ORNs) and their response is dampened by glutamate that is released from local neurons (LNs) and acts on GluClα receptors. (B) Illustration of the two-choice olfactory trap paradigm, where flies are given a choice to enter a trap containing an odor compound (red agar) or without odor (clear agar), and the number of trapped flies is analyzed after 24 hours. Preference of GluClα unedited and *wt* males towards apple juice (C) and ethanol (D) containing, n=20 ***P-value<0.0001; Mann-Whitney test. (E) Graphical depiction of Y maze assay in which flies can climb towards odors placed at the end of the arms and their amount on each side is used to calculate preference towards apple juice or ethanol. Preference of GluClα unedited and *wt* males towards apple juice (F) or ethanol (G) as analyzed by the Y maze assay. Unedited flies exhibit reduced attraction to apple juice and ethanol **P-value<0.01; T-test, n=20. *wt* flies exhibit preference to apple juice and ethanol that differs significantly than 0 (one sample t test; p<0.0001) whereas GluClα unedited do not differ than 0. (G) Schematics of the multiplex system which measures the fraction of time during which single flies move towards or away from acetic acid (I) and 2-butanone (J). Mean time of *wt* flies away from acetic differs or 2-butanone differs from zero (one sample t test; p value (two tailed) <0.0001) while unedited is no different from 0. Unedited flies show decreased aversion to acetic acid, n=35 ***P-value<0.0001; Mann Whitney test, and towards 2-butanone, n=40 ***P-value<0.0001; Mann-Whitney test.

Since attraction or aversion to odors requires intact motor capabilities, we first verified that *GluClα^unedited^* flies exhibit normal locomotor behaviors by analyzing various features of their motor actions using the FlyBowl tracking-based behavioral analysis system (Supp Fig. 2). Fruit flies use odors to navigate across complex environments and locate food sources^62^. Ethanol and amine volatiles emitted from rotting fruits attract *Drosophila* flies, who aggregate on fermenting fruits as a hub for feeding, mating, and egg laying^63–65^. To examine the attraction of *GluClα^unedited^* flies to appetitive odors, we used the simple two-choice olfactory trap assay, in which one of the traps contained an appetitive odor such as apple juice or ethanol, whereas the other trap contained plan agar as control (Fig. 2B). Fifty 3-day-old flies were introduced into chambers containing two Eppendorf-based traps with either 1% agar or agar supplemented with 10% apple juice and were left uninterrupted for 24 hours before the number of flies in each trap was counted. Comparing the preference of *wt* flies to that of *GluClα^unedited^* flies we discovered that while *wt* flies exhibited robust preference towards apple juice, *GluClα^unedited^* flies showed a marked reduction in the number of flies that entered the apple juice traps (Fig. 2C). The reduced preference towards appetitive odor was also apparent towards ethanol (Fig. 2D). Next, we tested the attraction towards each of these odors using a simple Y maze assay, in which flies can choose an arm containing at its end agar or agar with 10% apple juice or ethanol (Fig. 2E). Unlike the olfactory trap assay that measures the number of flies trapped in each of the traps over 24 hours, the Y maze is based on an immediate choice between the arms of the maze. Fifty 3-day-old *GluClα^unedited^* or *wt* flies were inserted at the entrance of each Y maze apparatus, and the flies were given 40 seconds to choose between the two arms before the vials were cupped. While *wt* flies show a strong preference to apple juice and ethanol-containing arms, the mean preference of *GluClα^unedited^* flies is close to zero (Fig. 2F, G), implying a random choice between the two arms. The decreased attraction of *GluClα^unedited^* flies to apple juice and ethanol in two independent assays suggests that editing of Il →e27 Val is necessary for proper olfactory responses toward appetitive odors.

Next, we analyzed the response of *GluClα^unedited^* flies to aversive odors using the multiplex system^66^. This single fly assay tracks temporal approaches towards an odor or avoidance from it, and as such overcome the inherent limitation of olfactory trap and Y-maze systems that measure behavioral choices of groups in which the choice of individuals may be influenced by those of other flies (Fig. 2H). We used the multiplex system to examine the responses of *GluClα^unedited^* flies to two aversive odors: acetic acid and 2-butanone. Examining the time spent avoiding each of the odors, showed that while the control (*wt*) flies had a negative value of close to -100, in agreement with robust avoidance, the average time depicted by GluClα^unedited^ flies was close to zero, suggesting that their movement within the chamber is random (Fig.2 I, J). These results demonstrate that editing of Ile^27^**→**Val in GluClα is necessary for proper chemotaxis.

### Recoding of Ile^27^ to Val is necessary for the expression of complex olfactory guided behaviors

Given the reduced response to appetitive and aversive odors, we next extended the analysis of GluClα^unedited^ flies to include behaviors that rely on olfactory cues, such as pheromone-based social interactions^67^. A central pheromone in this respect is the male-specific pheromone cVA that is known to induce male-male aggression and promote sexual receptivity in female flies^68^. Comparing aggression levels between pairs of GluClα^unedited^ flies to those of *wt* flies, revealed a dramatic reduction in the number of lunges exhibited by GluClα^unedited^ flies (Fig. 3A), as well as longer duration until they exhibited first lunge (latency to first lunge, Fig. 3B). The overall reduction in aggressive behavior suggests impaired perception of cVA. These striking results prompt us to examine another cVA-dependent behavior in female flies, where the presence of cVA emitted from courting male flies promotes receptivity^68–70^, measured by the time it takes from first courtship action until copulation (latency to copulation). For that, we paired GluClα^unedited^ or *wt* female flies with *wt* male flies and assayed their receptivity. First, we verified that *wt* male flies exhibit similar courtship patterns towards GluClα^unedited^ and *wt* females and discovered that the time it took male flies to initiate courtship action was similar towards the two genotypes (Fig. 3C). Next, we measured the time it took from initiation of courtship action by male flies and until the beginning of copulation. GluClα^unedited^ females exhibited significantly longer latency to copulate, which reflects reduced receptivity (Fig. 4D). The reduced receptivity agrees with impaired perception of cVA.

**Figure 3.**
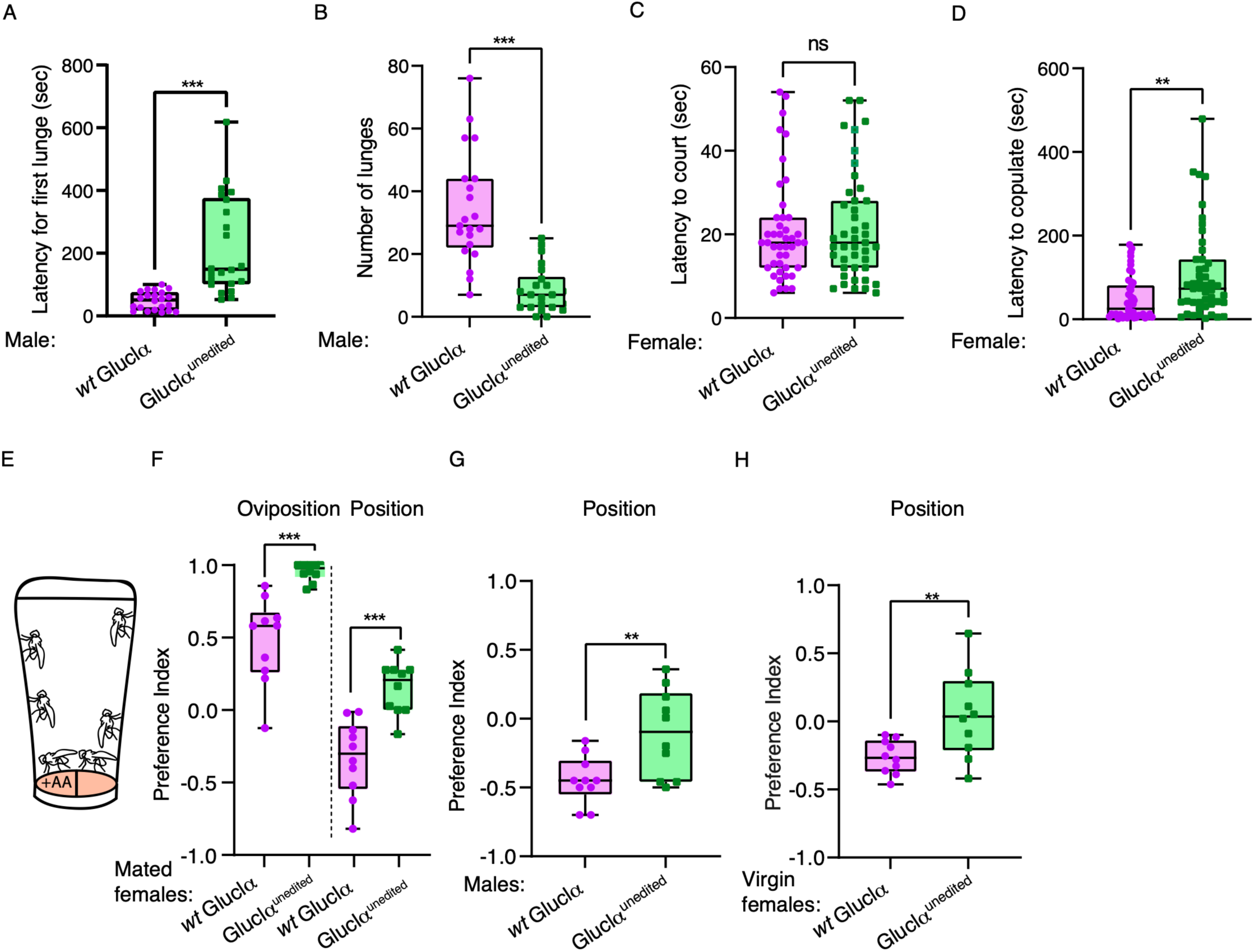
GluClα unedited flies exhibit impaired olfactory based behaviors. (A) GluClα unedited flies exhibit longer duration to first lunge during aggression assay. n=23, ***P-value<0.0001; Mann Whitney test. (B) GluClα unedited flies exhibit a reduced ratio of lunges. n=23, ***P-value<0.0001; Mann Whitney test. (C) *wt* male flies exhibit similar courtship behavior towards *wt* and GluClα unedited female flies, manifested by similar latency to first courtship action. n=50 P>0.05 Mann Whitney test. (D) GluClα unedited female flies exhibit reduced receptivity to courtship by *wt* male flies as measured by the latency to mate. take a similar amount of time to start courting mutant and wtcs females. n=50, **P-value<0.01; Mann-Whitney test. (E) Schematic illustration of two choice ovipositional and positional preference in egg laying on substrate with acetic acid. (F) Oviposition and position indices of mated GluClα unedited and *wt* females. GluClα unedited female exhibit enhanced oviposition index n=10 (***p value<0.0001 one sample t test) and reduced positional aversion to acetic acid (** p<0.005, t test, two-tailed) which significantly different from zero (*p<0.05, two-tailed t test). (G) GluClα unedited males show reduced positional aversion to substrate containing acetic acid n= 10 (test). H. Virgin GluClα7 unedited female exhibit reduced positional aversion to substrate containing acetic acid n=10 (test).

**Figure 4.**
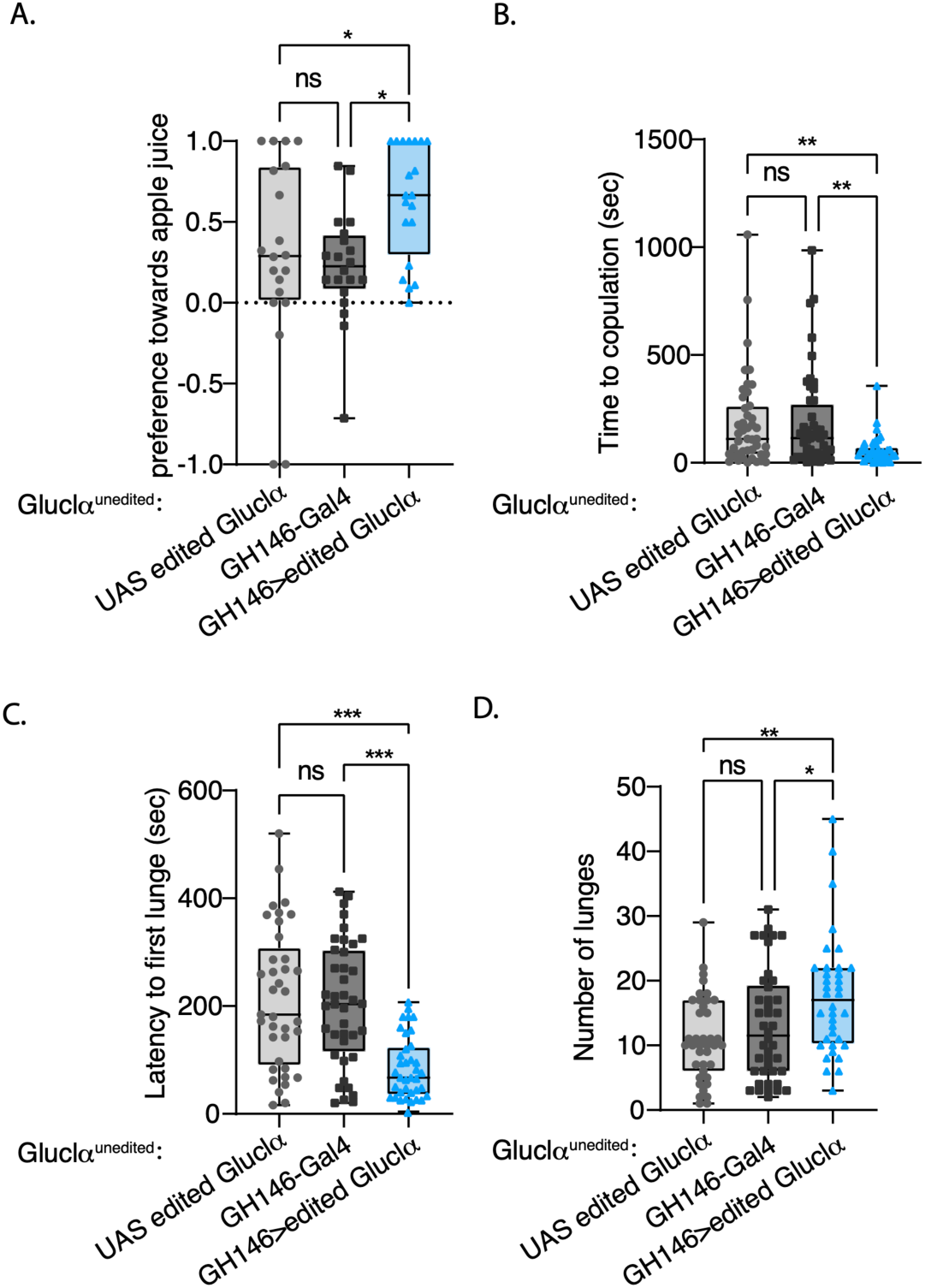
Expression of an edited channel in the PN in unedited GluClα flies manages to rescue the behavioral phenotype in behaviors involving smell sensing. (A) Flies expressing an edited channel in PN’s (bluish) show a significant increase in preference for the smell of apple juice in the olfactory trap test compared to the control flies (gray tones). *P-value<0.005; one-way Anova, n=20. (B) Females expressing an edited channel in PN’s show a significant decrease in the time it takes for them to start mating compared to the control females. **P-value<0.001; one-way Anova, n=45 (C) Males expressing an edited canal in PN’s show a significant decrease in the time it takes them to perform the first aggressive push ***P-value<0.0001; one-way Anova, n=40 (D) and an increase in the number of urges compared to the control flies. *P-value<0.005; one-way Anova, n=40. In all panels, colors are as in A.

We then explored the phenotype GluClα^unedited^ female flies in a two-choice egg-laying paradigm, in which females exhibit attraction to food containing acetic acid as an egg-laying substrate, and at the same time show aversion to staying on acetic acid-containing food (positional avoidance)^71^. The oviposition attraction to acetic acid is mediated by gustatory neurons, and the positional aversion by olfactory cues^71, 72^. The trade-off between oviposition preference and positional avoidance is an intriguing system to test the behavior of GluClα^unedited^ female flies. The experimental design consists of a simple apparatus in which mated females were allowed the choice to lay eggs on regular food or on food containing 5% acetic acid, and the number of eggs laid on both sides is counted after 3 hours. During this period, the position of females is monitored every 15 minutes to calculate positional preference/aversion towards acetic acid-containing substrate (Fig. 3E). As expected, *wt* female flies show a robust preference to lay eggs on food with acetic acid (positive oviposition preference values), while they avoid staying there (negative positional preference values, Fig. 3F), demonstrating the opposing motivations, between the need to lay eggs in a substrate containing acetic acid and their avoidance from its aversive odor. GluClα^unedited^ females, on the other hand, lost the positional aversion to acetic acid and instead showed a slight preference to be on acetic acid food, and accordingly showed a significantly higher preference to lay all their eggs on this substrate (Fig. 3F). Moreover, GluClα^unedited^ males and virgin female that lack the motivation to lay eggs, and therefore do not need to stay on acetic acid as a substrate for egg laying, do not show positional aversion to acetic acid, and spend on average equal time on both substrates (Fig. 3G, H). The results suggest that while the gustatory perception of acetic acid that is responsible for oviposition is normal in GluClα^unedited^ females, their olfactory perception of acetic acid is impaired, pointing to the importance of Ile^27^ recoding in the processing of olfactory information.

### The recoding of Il27→Val is necessary in projection neurons for the proper expression of olfactory-based behaviors

Considering the olfactory-related phenotypes observed in GluClα^unedited^ flies, and the fact that these flies express an unedited form of GluClα in all the cells that normally express GluClα, we next searched for the neurons in which the recoding of Ile27 to Val has a functional relevance. Since GluClα was previously shown to function in olfactory projection neurons (PNs)^40^, we tested whether expressing a fully edited version of GluClα in olfactory projection neurons of GluClα^unedited^ flies can rescue the olfactory phenotypes. To this end, we generated GluClα^unedited^ flies harboring the GH146 driver that target ∼90 of the 200 PNs in the AL^73^ and a UAS transgene that allows the expression of GluClα protein in which the six editing sites are in a fully edited state (the A nucleotide is mutated to G). We next profiled the behavior of GluClα^unedited^ flies expressing a fully edited version of GluClα in projection neurons using three behavioral paradigms. First, we examined their attraction to apple juice using an olfactory trap assay. While both genetic controls exhibit no preference for apple juice, the expression of edited GluClα in PNs led to a clear preference towards apple juice, indicating that the expression edited version can rescue the impairment in olfactory response to appetitive odors (Fig. 4A).

Furthermore, GluClα^unedited^ females harboring the edited form in PNs, exhibited a significant increase in their receptivity to courting males, as the latency to copulation was dramatically shorter than the genetic controls (Fig. 4B). Expressing the edited form in PNs rescued the reduced aggression observed in GluClα^unedited^ males, as the experimental males exhibited shorter duration to the expression of first lunge and higher number of lunges in comparison to genetic controls (Fig. 4C, D). Taken together, our results indicate that the loss of function phenotypes of GluClα^unedited^ flies can be partially rescued by the expression of edited version in PNs, suggesting that the recoding of Il27**→**Val has a functional role in PNs to facilitate proper olfactory responses.

### Recoding of I27V in GluClα shapes odor responses within the antenatal lobe

To strengthen the functional role of the edited form of GluClα in PNs, we next explored its effect on neuronal physiology, by analyzing the odor-evoked neuronal activity of PNs within the AL. It was previously shown that glutamate functions as an inhibitory transmitter that shapes PNs response to olfactory stimuli^40^. Glutamate that is secreted from local neurons (LNs), binds to GluClα on PNs and inhibits their activity, as knockdown of GluClα in PNs results in increased responses of certain glomeruli to isoamyl alcohol (IAA) and methyl salicilate^40^. Based on these findings, we expressed GCaMP6m in PNs in GluClα^unedited^ and *wt* flies and performed 2-photon functional Ca^2+^ imaging in response to isoamyl alcohol (IAA) and methyl salicylate.

We first validated that the GluClα^unedited^ flies harboring either the PNs driver or the GCaMP6m exhibit reduced attraction to appetitive odor compared to *wt* flies that harbor the same transgenes (Supp. Fig. 3). Next, we compared that odor-evoked activity across all glomeruli and found similar responses to both odors in GluClα^unedited^ and *wt* PNs (Fig. 5A). The overall similarity in the responses when examining all glomeruli prompt us to analyze the responses of single glomeruli across the antennal lobe including DM5, VA1v, VA2, VA3, VC1, VL2a, VL2p and VM2. Interestingly, while most tested glomeruli exhibited similar responses in GluClα^unedited^ and *wt* flies, the VA1v glomerulus exhibited significantly increased activity in GluClα^unedited^ flies in response to IAA and methyl salicylate (Fig. 5B). The enhanced responses of VA1v PNs to IAA and methyl salicylate in unedited flies, suggest that the recoding event of Ile^27^ in GluClα is necessary to dampen the activity of the PNs during the perception of these odors. These results support our behavioral finding, altogether indicating that the recoding of Ile^27^ is required for correct function of AL neurons in regulating olfactory based behavioral responses.

**Figure 5.**
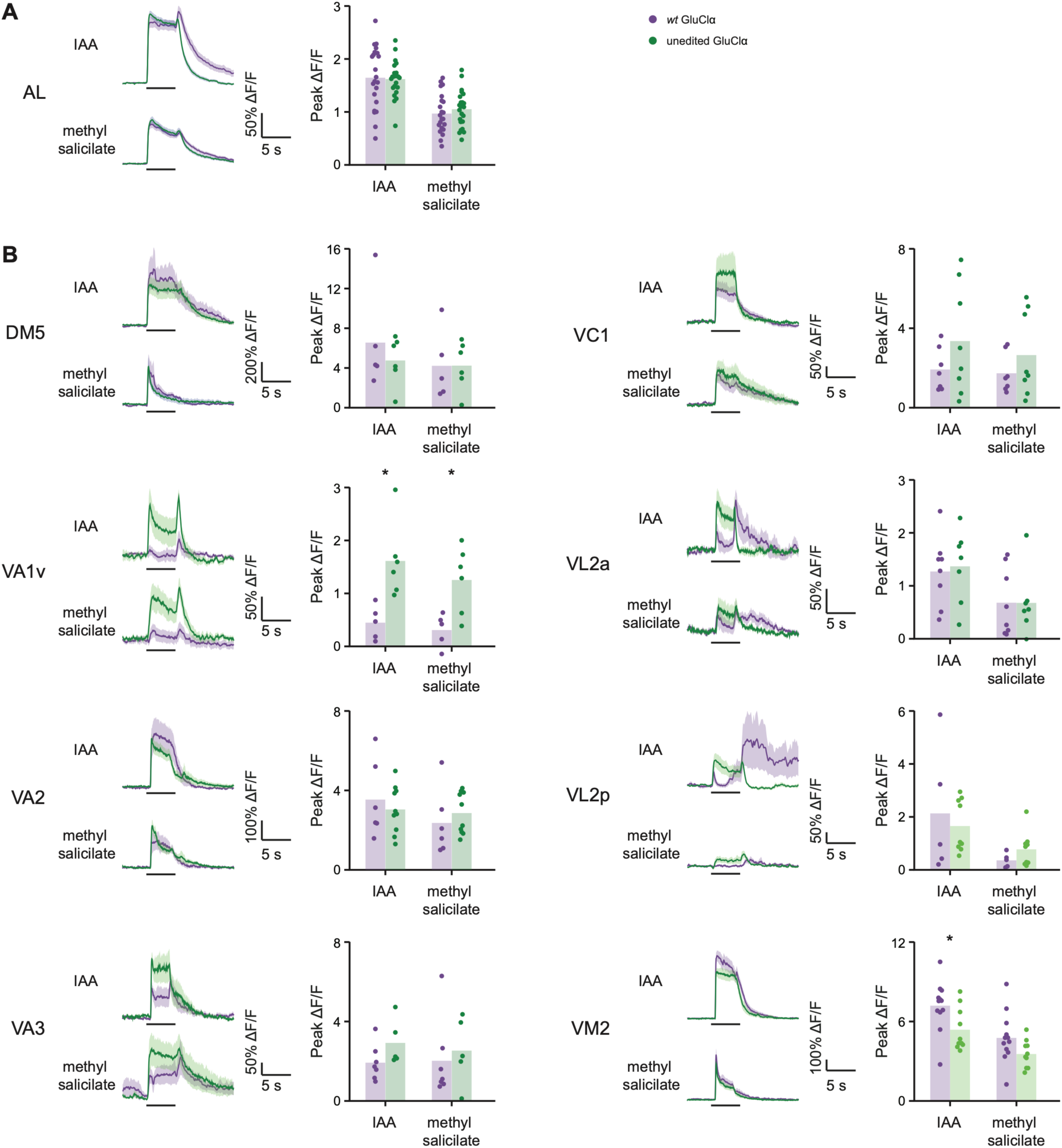
Unedited GluClα affects PN odor responses. (A) ΔF/F of GCaMP6f signal in the entire AL in control GluClα (purple) and unedited GluClα (green) flies, during presentation of odor pulses (isoamyl acetate, IAA, and methyl salicylate, horizontal lines). Data are mean (solid line) ± SEM (shaded area). No significant difference in odor responses was observed between flies with GluClα unedited background to the control flies with *wt* background. Paired t-test, n=5. (B) ΔF/F of GCaMP6f signals in eight different glomeruli to IAA and methyl salicylate in GluClα unedited flies (green) compared to wt flies (purple). A significant increase in odor response was observed in VA1v glomerulus for the two tested odors. * p-value<0.05; t-test, n=6.

## Discussion

A-to-I RNA editing has long been suggested as cellular machinery that can provide the proteomic diversity required for the intricate function of the nervous system, by shaping the spatial and temporal repertoire of expressed protein isoforms expressed in neurons^10, 74–77^. Here we dissected for the first time the physiological importance of an editing site in *Drosophila* and showed that a highly abundant and evolutionarily conserved recoding event that occurs within the extracellular domain of *GluClα* is necessary for proper olfactory responses in adult flies. Firstly, by ablating the editing of Ile^27^**→**Val, such that the endogenous channel harbors only the unedited version (Ile^27^ instead of Val) we were able to examine the contribution of this particular recoding event to its function in behaving animals. Secondly, by using a set of behavioral assays, we discovered that the edited isoform is necessary for the ability of flies to perceive appetitive and aversive odors, as well as for the expression of pheromone-based social behaviors. Furthermore, we mapped the spatial requirement of the edited isoform to olfactory projection neurons.

Consistent with a previous study^40^, our results indicate that the unedited isoform of *GluClα* affects odor responses in only one glomerulus out of the eight tested. Considering results from previous studies which found that silencing one or a few glomeruli had no effect on behavioral output^78^, it is unclear how the lack of *GluClα* editing in PNs has such a pronounced effect on behavior. It is important to note, that the affected glomerulus (VA1v) is sexually dimorphic^79–81^ receiving input from Or47b neurons known to respond to conspecific odors. PNs responding to food odors are known to be less specific than their cognate ORNs^82, 83^. In *GluClα^unedited^* flies, the VA1v glomerulus loses its specificity and responds to non-pheromone odors, suggesting that under normal conditions glutamatergic inhibition is required to maintain VA1v specificity. It is therefore possible that the loss of odor valence observed in unedited flies results from improper activation of pheromonal pathways during the perception of non-pheromonal odor signals, leading together to contradicting consequences. Whether this is indeed the case, is out of the scope of this manuscript. Furthermore, it would be intriguing to examine in future work whether *GluClα* has a role in controlling the activity in other pheromone related glomeruli.

The current case is not the first recoding event that is necessary for the proper function of a channel. The ablation of an editing site located within the mammalian excitatory ionotropic glutamate receptor subunit B (GluR-B) leads to early seizure-related death in mice^84^. While the ablation of Ile^27^ recoding in *GluClα* did not reduce viability or reproduction of flies under lab conditions, the impaired olfactory responses exhibited by unedited flies are expected to affect their fitness in natural environments, since flies heavily rely on their olfactory system to survive and reproduce.

The robust loss of function phenotype of the unedited form indicates that recoding of Ile^27^ to Val is necessary for the proper function of the channel, and the combination of high editing levels and evolutionary conservation are useful criteria when predicting which of the many editing events has the potential of having a function. This brings up the question of why evolution chose to maintain modulation of the sequence at the RNA level rather than inserting a mutation at the DNA level. The answer probably lies in the varying levels of its editing between different neuronal populations, suggesting that the ratio between edited and unedited forms of the channel or within subunits that construct the same channel may be spatially regulated to fine-tune its function. A relevant example in this respect is the spatial and temporal regulation of a recoding event that is found within the calmodulin-binding IQ-domain of the voltage-activated calcium channel (Cav1.3), whow ablation enhances learning^16, 17^. Further studies are required to test whether the editing repertoire of *GluClα* is regulated in response to different environmental conditions.

It is intriguing that a recoding event that is conservative in its nature (Ile**→**Val) and that is located close to the N terminal tip of the channel (five amino acids downstream to the putative signal peptide) has such a strong impact on function. As of this moment, there is no resolved 3-D structure of *Drosophila* GluClα, and 3-D structures of other Cys-loop channels lacked a portion of the N terminal segment that contains this particular residue, making it difficult to assess the molecular function of Ile^27^ recoding to Val. Still, there are striking examples of the way by which a similar recoding event (Ile^400^**→**Val) that is located at the intracellular cavity of the potassium channel Kv1.1 strongly modifies the inactivation kinetics, where the edited form of the channel recovers from inactivation 20 times faster than the unedited form^85^. The faster recovery is achieved due to reduced hydrophobic interaction between the tip of the inactivation gate and the Val residue that is found within the intracellular cavity^85^. While the Ile^27^-containing segment of the *Drosophila* GluClα subunit is missing in the various 3-D structures of anionic Cys-loop receptors (Supporting Figure. 1), its location is clearly a few amino acids upstream to the N-terminal α helix (α1) that is typical of Cys-loop receotors (Suup. Fig. 1). Helix α1 lies in close proximity to a conserved amino-acid cluster that belongs to the adjacent subunit (e.g., Fig. 6A-C, for cluster sequence see Supp. Fig 1). This close proximity allows for van der Waals interactions^86^ (Fig. 6C), and in some cases hydrogen bonding^87^ (e.g., PDB 6PLR^87^), between entities of helix α-helix and amino acids belonging to the aforementioned cluster in the adjacent subunit. One may therefore envision that in the *Drosophila* GluClα receptor the N-terminus and the editing site itself might be close, at an interaction distance, to the neighboring subunit. Since inter-subunit interactions often take part in the function of Cys-loop receptors, further studies are necessary to assess how the recoding of this position affects the biophysical properties of the *Drosophila* GluClα receptor.

**Figure 6.**
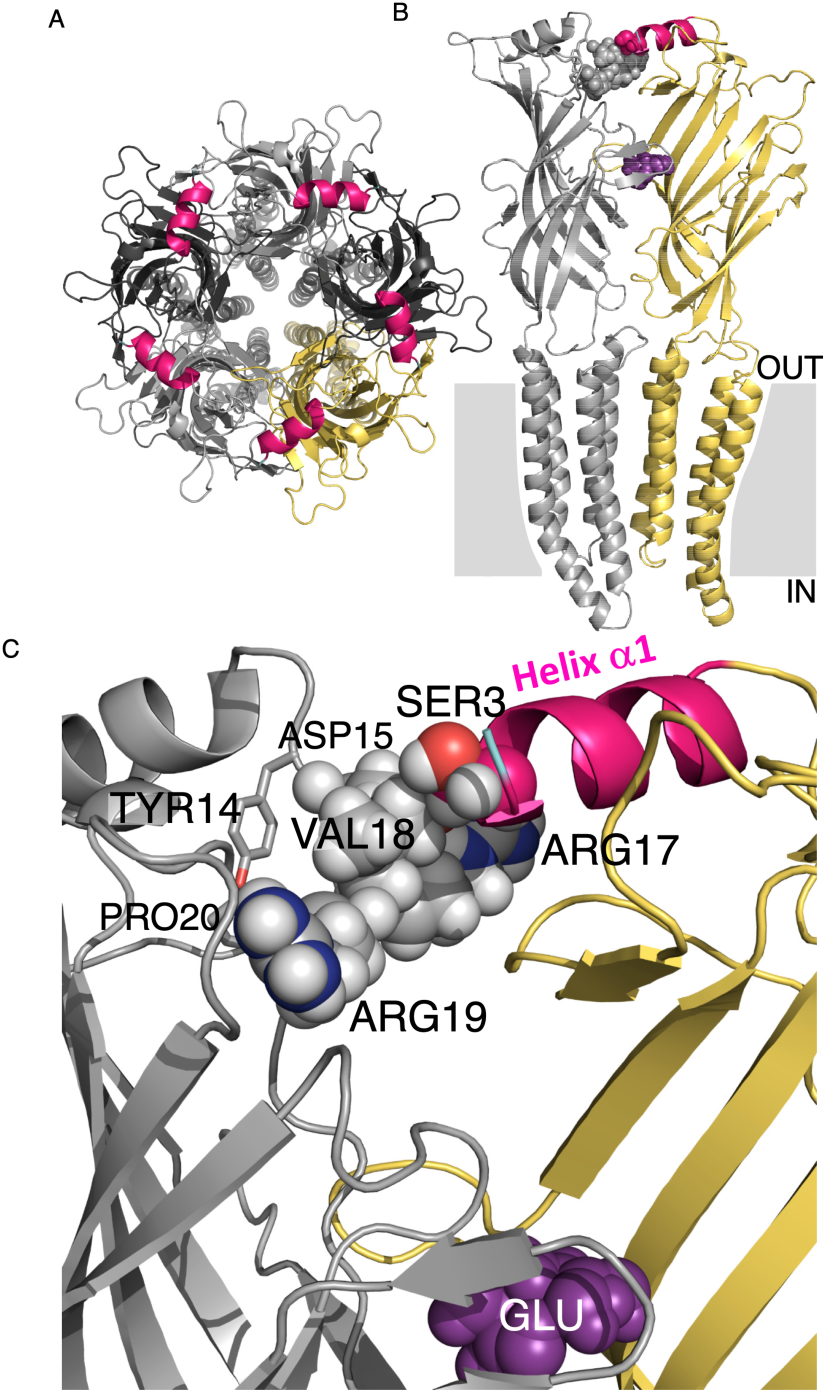
Structural characteristics of GluCl receptors. (A) Top view of the *C. elegans* GluClα receptor (GluClαcrystR; Protein Data Bank (PDB) ID code 3RIF) showing five identical subunits, which are colored differently to highlight the intersubunit interfaces. Alpha-helix α1 is colored in pink in all five subunits. (B) Two adjacent subunits (colored in gray and yellow) of the GluClαcrystR are viewed from the side. The membrane region is illustrated as a gray rectangle interrupted by the transmembrane helices (OUT, extracellular; IN, intracellular). The N-terminal tip of α-helix α1 (pink), which belongs to the yellow-colored subunit, interacts with the adjacent subunit (gray spheres), as detailed in panel C. The neurotransmitter glutamate (purple spheres) is bound at the intersubunit interface. (C) Space-filling models of the side chains of SER3 (belongs to Helix α1) and VAL18 (belongs to the conserved cluster shown in Supporting Figure 1) indicate van der Waals interactions between these two amino acids (3.9 Å center-to-center distance between O^ψ^ of SER3 and C^ψ^^2^ of VAL18). Side chains of the other amino acids belonging to the conserved cluster are shown as spheres or sticks. Carbon, oxygen, nitrogen and hydrogen atoms are colored in pink or gray, red, blue and white, respectively. GLU (the neurotransmitter) is shown in purple spheres.

## Materials and Methods

### Analyzing evolutionary conservation of GluClα editing sites across *Drosophila* species

The analysis is based on an existing data set from Ramaswami et al.,^88^ that quantified RNA editing at 605 loci using a multiplex microfluidic PCR with deep sequencing across 131 Drosophila strains (n=2/strain). The study used multiplex PCR primers to amplify 605 loci that include many known RNA editing events. PCR products of each sample were then subjected to a 15-cycle barcode PCR and pooled together. The library was sequenced using Illumina HiSeq with 101 bp paired end reads. Paired-end reads were combined and mapped onto the genome (dm3) using BWA samse allowing 9 mismatches per read. The sequencing reads were aligned to a combination of the reference genome and 100 bp exonic sequences surrounding known splicing junctions from available gene models (obtained from the UCSC genome browser). We quantified editing levels of known *D. melanogaster* RNA editing sites by taking the fraction of reads containing a ‘G’ nucleotide at that position. For editing level quantification, sites covered by ≥50 mmPCR-seq reads were used. For each strain, we excluded editing sites where the measured editing levels in the two biological replicates differed by >20% (see Fig. 1b). Custom scripts used to process data are available upon request. In this study we compared the relative editing levels of 6 editing sites in GluClα.

### Analysis of GluClα editing levels across the fly brain

We used an existing dataset^47^ and used the following analysis pipeline: STAR (v2.4.2) (1) (--twopassMode Basic) was used to map paired-end mmPCR-seq reads and single-end RNA-seq reads to the dm6 genome as described above. We then used the Samtools mpileup function to determine base calls from uniquely mapped reads at known and novel editing sites, and calculated editing levels as number of G reads divided by the total of both A and G reads at a site. For mmPCR-seq, we required each replicate to have 100X coverage and we removed sites that were not within 20% editing between replicates, as done previously^88^. Final mmPCR editing levels were determined after down sampling coverage to 200 reads for statistical analysis.

### Fly lines

All *Drosophila melanogaster* fly lines used in this study were kept at 25C°, ∼50% humidity, light/dark of 12:12 hours, and maintained on cornmeal, yeast, molasses, and agar medium. Most fly lines were backcrossed to a Canton S background, or if mentioned were maintained on w1118 background. *wt* GluClα and unedited GluClα were used for behavioral assays. The genotype of GluClα unedited flies was validated in each experiment using PCR analysis (see below). GH146-Gal4>UAS GCaMP6 harboring *wt* or unedited version of GluClα were used for the Ca^2+^ imaging experiments. UAS edited GluClα flies were a generous gift from Liam Keegan.

### Generation of GluClα unedited flies

Ablation of Il27 editing site was achieved by deletion of the of intronic ECS region: chr3R: 15578744-15579171 using CRISPR/Cas9 and the following gRNAs to induce double-stranded cuts to the DNA, on each side of the ECS: TCTAAACCCTAGATATACGCTGG and TATAGTATGTGACTTTGCCTGGG. In addition, we made synonymous mutations to the sgRNA target region or Protospacer Adjacent Motif (PAM) to prevent recutting by the Cas9/CRISPR system. The genotype of the GluClα unedited flies was validated by performing simple PCR analysis using the following primers:

Forward: TTGTCTCCCGCTCCACTTAC Reverse: TTGGGCAATTTGAAAGTCGAAA

### Western blot analysis

The relative expression levels of *wt* and the unedited version of GluClα was assayed using western blots. Lysate was extracted from 20 heads of flies by RIPA buffer containing proteinase inhibitor and heated with sample buffer. 50 mg Protein was loaded in a 10% agarose gel in buffer run with the protein marker: And after an hour’s transfer, the membrane was blocked with 5% milk and exposed o.n. to a primary antibody against glucl and flag produced from rabbits and exposed for anti-rabbit secondary antibody. For control we also used an antibody against actin which was produced from mice at a concentration of 1:1000.

### Behavioral experiments

All behavioral experiments were performed about an hour after the lights on.

#### FlyBowl

Male flies were collected upon eclosion and aged in groups of 10 flies/vial for four days before test. 10 flues of each genotype were inserted in groups of 10 into Fly Bowl arenas^89^, and their behavior was recorded for 30 minutes and analyzed using CTRAX, FixTrax^90^ and JAABA^89^. For kinetic features, scripts were written in MATLAB to use the JAABA code to generate the statistical features as specified in Kabra et al,^89^. Time series graphs (per frame) were created using JAABA Plot^89^. Quantification of complex behaviors was done using JAABA Classifiers^89^ to identify specific behaviors: Walk, Stop, Turn, Approach, Touch, Chase, Chain, Song, Social Clustering, and Grooming. Bar graphs were created using JAABA Plot^89^. Each feature of the FlyBwol experiment was standardized according to all values calculated in our experiments for that feature to generate a z-score. Scatter plots were created using R.

#### Olfactory Trap assay

Traps consisting of an eppendorf tube connected to a 200 µl pippet tip was fitted to the trap to prevent flies from being able to escape the odor trap after entering it. Each trap is filled with 500 µl of a 1% agar supplemented with 10% odor apple juice^65^ or ethanol or water as a control. Each setup contains two traps (with odor and control) that are glued to a 90mm x 15mm Petri dish. The 2 traps are placed in opposite directions. 50 3-day-old flies are introduced into each such experimental system. The flies can be attracted to one of the traps based on smell alone, since they cannot make physical contact with the substrate to make the decision. To increase the motivation to choose one of the traps, the test was done in the absence of food. The experimental system is wrapped in aluminum foil to prevent the penetration of light that could disrupt their decision to enter one of the traps based solely on smell. The number of flies in each trap is counted after 24 hours to calculate preference index: Preference Index (PI) =(# flies that entered the trap with the tested odor -# flies that entered the trap that does not contain an odor source)/(total #flies).

#### Immediate odor preference using Y-maze

The behavioral setup is composed of two vials containing agar with and without odors that are connected to the Y-shaped adapter. Vial containing 50 3-day-old flies is placed at the entrance of the adapter, and the flies are allowed to choose one of the arms. After 40 seconds, the upper vials are closed and the number of flies in each is counted. Preference Index is calculated by as follows: Preference Index (PI) = (# flies in test arm-# flies in control arm)/(total #flies).

#### Single-fly assay for odor preference

Behavioral experiments were performed in a custom-built, fully automated apparatus as described in detail^66^. Single flies were introduced into clear polycarbonate chambers (length 50 mm, width 5 mm, height 1.3 mm) and were exposed to air containing odor flow. Odors were prepared at 10-fold dilution in mineral oil. Fresh odors were prepared daily. Air or odor streams from the two halves of the chamber converged at a central choice zone. The location of the flies was recorded and tracked. A fly’s preference was calculated as the percentage of time that it spent on one side of the chamber.

#### Two-Choice Oviposition and Positional Assays

The 2-choice egg laying setup contains a standard fly bottle, with the base cut off and replaced with a 60-mm Petri dish lid. Half of the Petri dish lid contains solid food substrate with 5% acetic acid, and the other half contains food supplemented with equal volume of water. Two-choice dishes were made by dividing a 35-mm Petri dish lid with a razor blade and pouring 2 samples of food-substrate into each half. For each test, 15–20 recently mated females were introduced into each two-choice apparatus allowed to sample and lay eggs for 3 h. Oviposition preference was determined by counting the number of eggs on each half of the 2-choice dish (oviposition index= # eggs laid on acetic acid containing food-# eggs laid on control food / #total eggs laid). For positional preference, the number of flies on each half of the dish was counted every 15 minutes along the entire duration of the experiment (3 h) (position index = average # flies on acetic acid half - average # flies on control food half / Totla # flies).

#### Courtship and mating tests

Male flies were collected upon eclosion and aged in groups of 10 flies/vial for four days before test. Courtship arenas were placed in behavior chambers, under controlled temperature and humidity (25°C, 70% humidity). Behavior was recorded for one hour from the introduction of male and female pairs using Point-Grey Flea3 cameras (1080×720 pixels at 30 fps). Latency to copulate was quantified for each pair as total time, starting from first wing vibration the male exhibited and ending in successful copulation.

#### Aggression

4 days old pairs of Single housed male flies (*wt*, GluClα or when indicated GH146>UAS edited GluClα and genetic controls) were put into round aggression arenas (about 0.08 cm3 in volume). A mixture of agarose and apple juice (1% agarose, 50% apple juice) was inserted into arenas to enhance aggressive behavior. Experiments were performed at a similar time of day (Lights ON + 1h). Flies’ behavior was recorded for 30 min with Point-Grey Flea3 (1080×720 pixels at 60 fps). Aggressive behavior was later quantified by counting the number of lunges for each pair, and latency as the time from start of experiment to first lunge for each pair.

#### Functional imaging

Flies used for functional imaging were raised on cornmeal agar under a 12 h light/12 h dark cycle at 25 °C. Imaging was done as previously described (Manoim et al 2022, Rozenfeld et al 2019, Israel et al 2022). Briefly, flies were anesthetized on ice then a single fly was moved to a custom-built chamber and fixed to aluminum foil using wax. Cuticle and trachea in the required area were removed, and the exposed brain was superfused with carbonated solution as described above. Odors (purest level available) were obtained from Sigma-Aldrich (Rehovot, Israel). Odor flow of 0.4 l/min (10-1 dilution) was combined with a carrier air stream of 0.4 l/min using mass-flow controllers (Sensirion) and software-controlled solenoid valves (The Lee Company). This resulted in a final odor dilution of 5X10-2 delivered to the fly. Odor flow was delivered through a 1/16 inch ultra-chemical-resistant Versilon PVC tubing (Saint-Gobain, NJ, USA) placed 5 mm from the fly’s antenna. Functional imaging was performed using a two-photon laser-scanning microscope (DF-Scope installed on an Olympus BX51WI microscope). Fluorescence was excited by a Ti-Sapphire laser (Mai Tai HP DS, 100 fs pulses) centered at 910 nm, attenuated by a Pockels cell (Conoptics) and coupled to a galvo-resonant scanner. Excitation light was focused by a 20X, 1.0 NA objective (Olympus XLUMPLFLN20XW), and emitted photons were detected by GaAsP photomultiplier tubes (Hamamatsu Photonics, H10770PA-40SEL), whose currents were amplified (Hamamatsu HC-130-INV) and transferred to the imaging computer (MScan 2.3.01). All imaging experiments were acquired at 30 Hz.

### Author Contributions and Notes

H.Z, G.S.O, J.B.L and M.P designed research, H.Z, P.D and E.R performed research, D.G and Y.P performed computational analysis, F.M.L, H.Z and E.R analyzed data; and H.Z, M.P, Y.P and G.S.O. wrote the paper.

The authors declare no conflict of interest.

This article contains supporting information online.

## Acknowledgments

We thank all members of the Shohat-Ophir lab for fruitful discussions and technical support. We would also like to thank Jennifer I. C. Benichou for statistical consultation. This work was supported by The U.S.-Israel Binational Science Foundation (BSF) Grant 2019091and the Israel Science Foundation Grant 174/19.

---

**Supplementary Figure 1.**
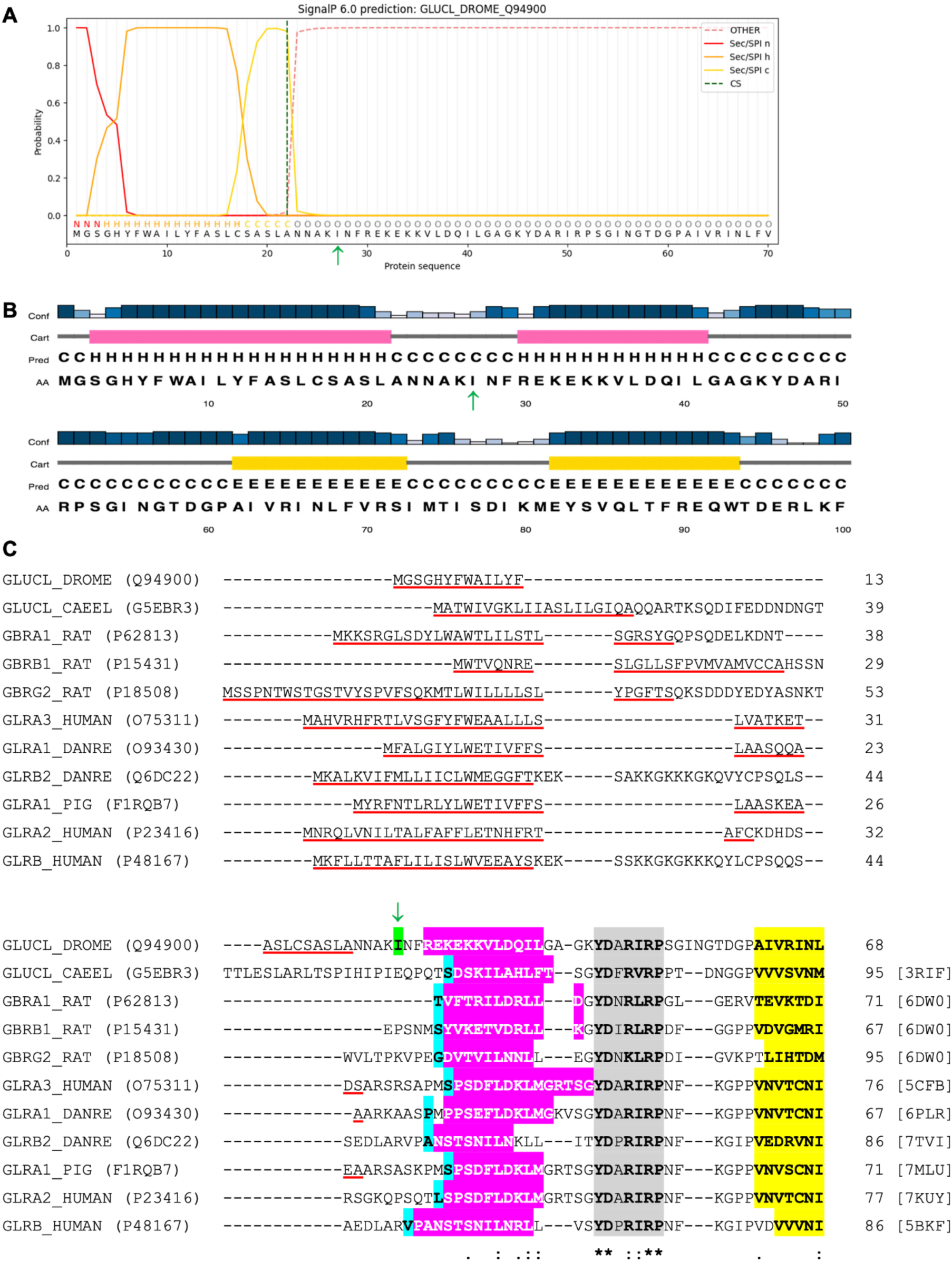
Location of the I27V editing site in the *Drosophila* GluClα subunit (UniProt ID Q94900). (A) SignalP 6.0 prediction showing that the probability of the presence of a signal-peptide cleavage site (CS) between positions 22 and 23 is *P* = 0.98. The Secretory (Sec) signal peptide n-region (Sec/SPI n) corresponds to the n-terminal region of the signal peptide and is labeled as N (above the amino acid sequence). Sec/SPI h corresponds to the center hydrophobic region of the signal peptide and is labeled as H. Sec/SPI c corresponds to the c-terminal region of the signal peptide and is labeled as C. OTHER corresponds to a sequence other than a signal peptide and is labeled as O. I27 is indicated by a green arrow. (B) PSIPRED 4.0 secondary structure prediction for the *Drosophila* GluClα immature protein. The full-length protein sequence was submitted for the prediction, whereas only the first 100 amino acids are shown. The protein secondary structure prediction is based on position-specific scoring matrices (PSSMs). Helices (H), strands (E) and coils (C) are indicated in pink, yellow and gray colors, respectively. Conf, confidence of prediction with the highest confidence in dark blue. AA, the target sequence. I27 is indicated by a green arrow. Note that transmembrane (TM) protein topology prediction based on support vector machines (SVM) (MEMSAT-SVM in the PSIPRED server) predicted amino acids 2-20 of the immature protein to be a signal peptide a-helix (not shown). (C) Iterative multiple sequence alignment between the *Drosophila* GluClα subunit (UniProt ID Q94900) and various subunits of anionic Cys-loop receptors whose three-dimensional (3-D) structure was determined at high resolution by X-ray crystallography or cryo-electron microscopy. For clarity, merely an N-terminal portion of the sequences is presented. The UniProt names and identifiers are indicated on the left side of the sequences. The PDB ID codes that correspond to the 3-D structures of the recombinant Cys-loop anionic receptors are indicated in square parentheses on the right side of the sequences. Signal peptides are underlined in red according to the UniProt Knowledgebase (UniProtKB). The edited amino acid (I27) is boxed in green and indicated by a green arrow. The first amino acid of each 3-D structure is boxed in cyan. Helix α1 is shown in white letters on a pink background. The highly conserved amino-acid cluster mentioned in the manuscript text is indicated by a gray background. The beginning of the first beta strand (b1) is indicated by a yellow background. Asterisk indicates positions which have a single, fully conserved residue. Colon indicates conservation between groups of strongly similar properties. Period indicates conservation between groups of weakly similar properties.

**Supplementary Figure 2.**
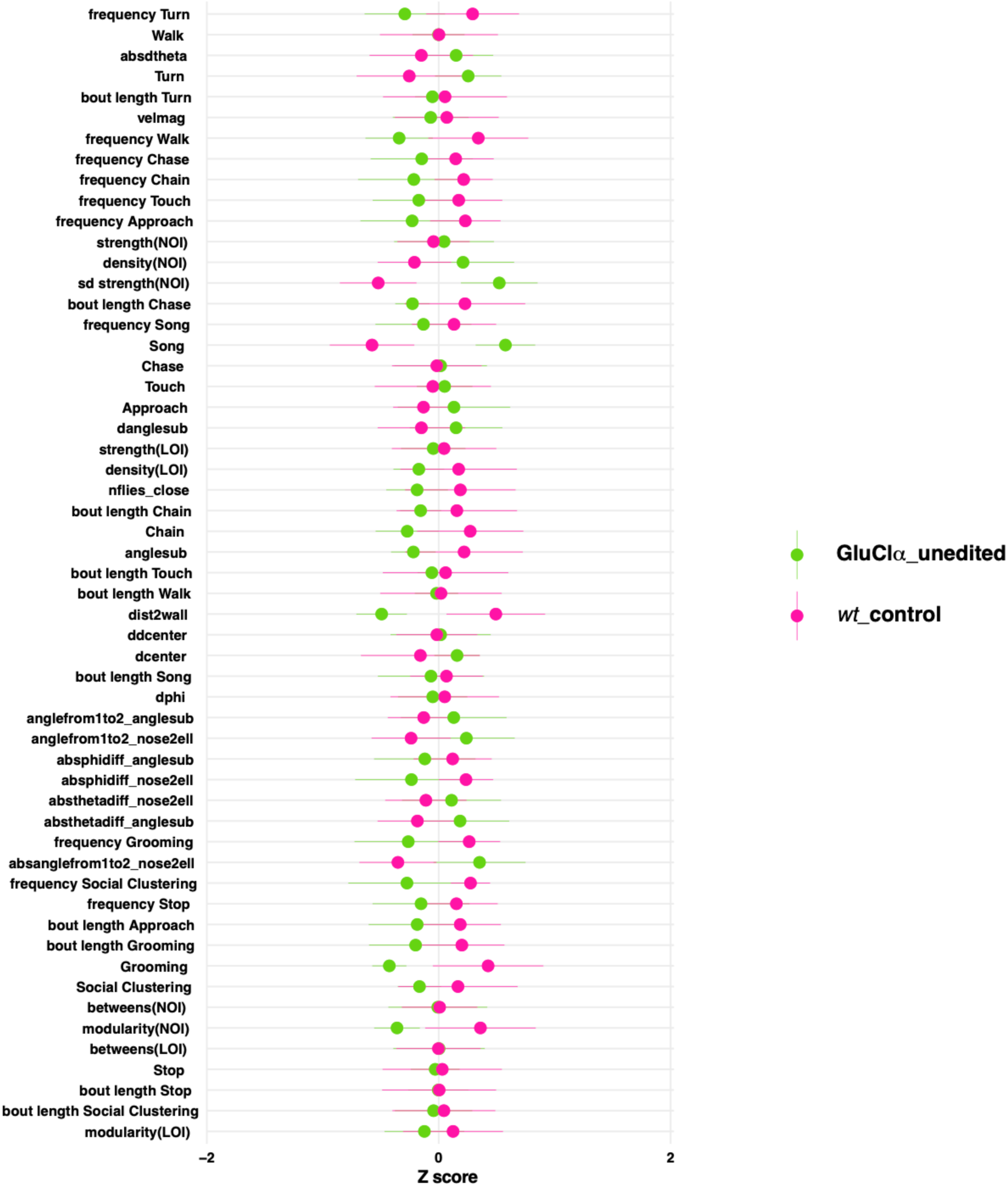
GluClα unedited flies show normal kinetic and social parameters in the FlyBowl system. Data are represented as normalized Z scores of 60 behavioral parameters, n = 7. No significant differences between GluClα unedited and *wt* male flies. Statistical analysis was determined by one-way ANOVA followed by Tukey’s range test for experiments that were distributed normally, and by Kruskal–Wallis test followed by Wilcoxon signed-rank test for experiments that were not distributed normally. FDR correction was used for multiple tests.

**Supplementary Figure 3.**
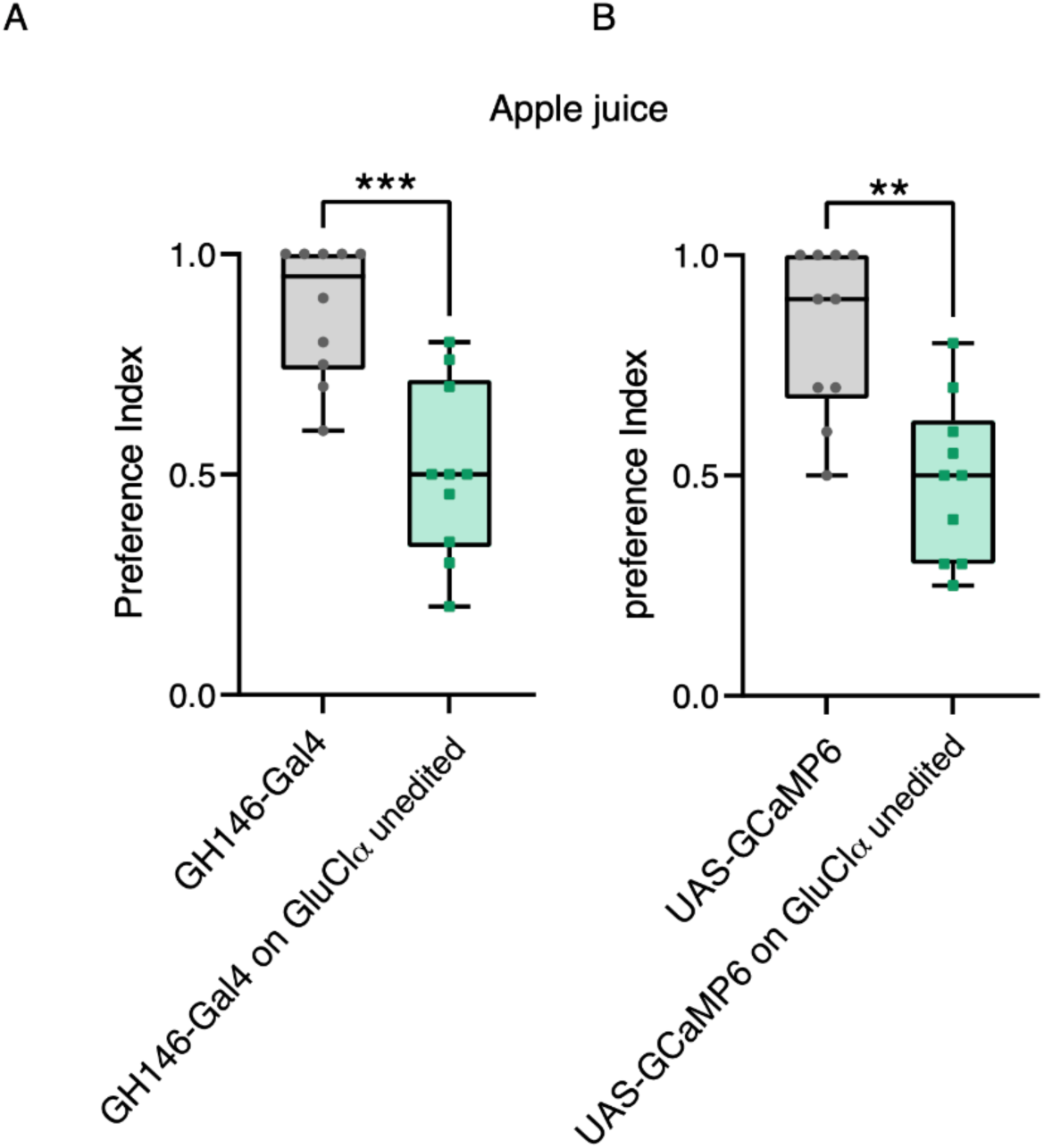
Validation strains for calcium imaging. (A) Preference of GH146-Gal4 lines and GH146-Gal4 on GluClα unedited males towards apple juice. n=10 ***P-value<0.0001; Mann-Whitney test. (B) Preference of UAS-GCaMP6 lines and UAS-GCaMP6 on GluClα unedited males towards apple juice. n=10 ***P-value<0.001; Mann-Whitney test.

Supp table 1: https://www.dropbox.com/s/zejjhdw2te7yalb/Supp%20table%201.csv?dl=0

Supp table 2a: https://www.dropbox.com/s/0tunrguqnyy55zd/Supp%20table%202a.csv?dl=0

Supp table 2b: https://www.dropbox.com/s/f1ggafojvip1a4x/Supp%20table%202b.csv?dl=0

Supp table 3: https://www.dropbox.com/s/9y12u483baahsi0/Supp%20table_3.csv?dl=0

## References

1. Eisenberg, E., and Levanon, E.Y. (2018). A-to-I RNA editing - immune protector and transcriptome diversifier. Nat. Rev. Genet. 19, 473–490.

2. Bass, B.L. (2002). RNA editing by adenosine deaminases that act on RNA. Annu. Rev. Biochem. 71, 817–846.

3. Roth, S.H., Levanon, E.Y., and Eisenberg, E. (2019). Genome-wide quantification of ADAR adenosine-to-inosine RNA editing activity. Nat. Methods 16, 1131– 1138.

4. Reautschnig, P., Wahn, N., Wettengel, J., Schulz, A.E., Latifi, N., Vogel, P., Kang, T.-W., Pfeiffer, L.S., Zarges, C., Naumann, U., et al. (2022). CLUSTER guide RNAs enable precise and efficient RNA editing with endogenous ADAR enzymes in vivo. Nat. Biotechnol. 40, 759–768.

5. George, C.X., Ramaswami, G., Li, J.B., and Samuel, C.E. (2016). Editing of Cellular Self-RNAs by Adenosine Deaminase ADAR1 Suppresses Innate Immune Stress Responses. J. Biol. Chem. 291, 6158– 6168.

6. Liddicoat, B.J., Piskol, R., Chalk, A.M., Ramaswami, G., Higuchi, M., Hartner, J.C., Li, J.B., Seeburg, P.H., and Walkley, C.R. (2015). RNA editing by ADAR1 prevents MDA5 sensing of endogenous dsRNA as nonself. Science 349, 1115–1120.

7. Duan, Y., Dou, S., Luo, S., Zhang, H., and Lu, J. (2017). Adaptation of A-to-I RNA editing in Drosophila. PLoS Genet. 13, e1006648.

8. Li, X., Overton, I.M., Baines, R.A., Keegan, L.P., and O’Connell, M.A. (2014). The ADAR RNA editing enzyme controls neuronal excitability in Drosophila melanogaster. Nucleic Acids Res. 42, 1139–1151.

9. Grassi, L., Leoni, G., and Tramontano, A. (2015). RNA editing differently affects protein-coding genes in D. melanogaster and H. sapiens. Scientific Reports 5. 10.1038/srep11550.

10. Garrett, S., and Rosenthal, J.J.C. (2012). RNA editing underlies temperature adaptation in K+ channels from polar octopuses. Science 335, 848–851.

11. Stapleton, M., Carlson, J.W., and Celniker, S.E. (2006). RNA editing in *Drosophila melanogaster*: New targets and functional consequences. RNA 12, 1922–1932. 10.1261/rna.254306.

12. Ryan, M.Y., Maloney, R., Reenan, R., and Horn, R. (2008). Characterization of five RNA editing sites in Shab potassium channels. Channels 2, 202–209.

13. Higuchi, M., Maas, S., Single, F.N., Hartner, J., Rozov, A., Burnashev, N., Feldmeyer, D., Sprengel, R., and Seeburg, P.H. (2000). Point mutation in an AMPA receptor gene rescues lethality in mice deficient in the RNA-editing enzyme ADAR2. Nature 406, 78–81. 10.1038/35017558.

14. Gaisler-Salomon, I., Kravitz, E., Feiler, Y., Safran, M., Biegon, A., Amariglio, N., and Rechavi, G. (2014). Hippocampus-specific deficiency in RNA editing of GluA2 in Alzheimer’s disease. Neurobiol. Aging 35, 1785–1791.

15. Venø, M.T., Bramsen, J.B., Bendixen, C., Panitz, F., Holm, I.E., Öhman, M., and Kjems, J. (2012). Spatio-temporal regulation of ADAR editing during development in porcine neural tissues. RNA Biol. 9, 1054–1065.

16. Huang, H., Tan, B.Z., Shen, Y., Tao, J., Jiang, F., Sung, Y.Y., Ng, C.K., Raida, M., Köhr, G., Higuchi, M., et al. (2012). RNA editing of the IQ domain in Ca(v)1.3 channels modulates their Ca^2+^-dependent inactivation. Neuron 73, 304–316.

17. Zhai, J., Navakkode, S., Yeow, S.Q.Z., Krishna-K, K., Liang, M.C., Koh, J.H., Wong, R.X., Yu, W.P., Sajikumar, S., Huang, H., et al. (2022). Loss of Ca1.3 RNA editing enhances mouse hippocampal plasticity, learning, and memory. Proc. Natl. Acad. Sci. U. S. A. 119, e2203883119.

18. Bazak, L., Haviv, A., Barak, M., Jacob-Hirsch, J., Deng, P., Zhang, R., Isaacs, F.J., Rechavi, G., Li, J.B., Eisenberg, E., et al. (2014). A-to-I RNA editing occurs at over a hundred million genomic sites, located in a majority of human genes. Genome Res. 24, 365–376.

19. Peng, Z., Cheng, Y., Tan, B.C.-M., Kang, L., Tian, Z., Zhu, Y., Zhang, W., Liang, Y., Hu, X., Tan, X., et al. (2012). Comprehensive analysis of RNA-Seq data reveals extensive RNA editing in a human transcriptome. Nat. Biotechnol. 30, 253–260.

20. Ramaswami, G., Lin, W., Piskol, R., Tan, M.H., Davis, C., and Li, J.B. (2012). Accurate identification of human Alu and non-Alu RNA editing sites. Nat. Methods 9, 579–581.

21. Porath, H.T., Schaffer, A.A., Kaniewska, P., Alon, S., Eisenberg, E., Rosenthal, J., Levanon, E.Y., and Levy, O. (2017). A-to-I RNA Editing in the Earliest-Diverging Eumetazoan Phyla. Molecular Biology and Evolution 34, 1890–1901. 10.1093/molbev/msx125.

22. St Laurent, G., Tackett, M.R., Nechkin, S., Shtokalo, D., Antonets, D., Savva, Y.A., Maloney, R., Kapranov, P., Lawrence, C.E., and Reenan, R.A. (2013). Genome-wide analysis of A-to-I RNA editing by single-molecule sequencing in Drosophila. Nat. Struct. Mol. Biol. 20, 1333–1339.

23. Graveley, B.R., Brooks, A.N., Carlson, J.W., Duff, M.O., Landolin, J.M., Yang, L., Artieri, C.G., van Baren, M.J., Boley, N., Booth, B.W., et al. (2011). The developmental transcriptome of Drosophila melanogaster. Nature 471, 473–479.

24. Sinigaglia, K., Wiatrek, D., Khan, A., Michalik, D., Sambrani, N., Sedmík, J., Vukić, D., O’Connell, M.A., and Keegan, L.P. (2019). ADAR RNA editing in innate immune response phasing, in circadian clocks and in sleep. Biochim. Biophys. Acta Gene Regul. Mech. 1862, 356–369.

25. Deng, P., Khan, A., Jacobson, D., Sambrani, N., McGurk, L., Li, X., Jayasree, A., Hejatko, J., Shohat-Ophir, G., O’Connell, M.A., et al. (2020). Adar RNA editing-dependent and -independent effects are required for brain and innate immune functions in Drosophila. Nat. Commun. 11, 1580.

26. Palladino, M.J., Keegan, L.P., O’Connell, M.A., and Reenan, R.A. (2000). A-to-I pre-mRNA editing in Drosophila is primarily involved in adult nervous system function and integrity. Cell 102, 437–449.

27. Savva, Y.A., Jepson, J.E.C., Sahin, A., Sugden, A.U., Dorsky, J.S., Alpert, L., Lawrence, C., and Reenan, R.A. (2012). Auto-regulatory RNA editing fine-tunes mRNA re-coding and complex behaviour in Drosophila. Nat. Commun. 3, 790.

28. Jepson, J.E.C., Savva, Y.A., Yokose, C., Sugden, A.U., Sahin, A., and Reenan, R.A. (2011). Engineered alterations in RNA editing modulate complex behavior in Drosophila: regulatory diversity of adenosine deaminase acting on RNA (ADAR) targets. J. Biol. Chem. 286, 8325–8337.

29. Jepson, J.E., and Reenan, R.A. (2010). Unraveling pleiotropic functions of A-to-I RNA editing in Drosophila. Fly 4, 154–158.

30. Keegan, L.P., Brindle, J., Gallo, A., Leroy, A., Reenan, R.A., and O’Connell, M.A. (2005). Tuning of RNA editing by ADAR is required in Drosophila. EMBO J. 24, 2183–2193.

31. Jepson, J.E.C., and Reenan, R.A. (2009). Adenosine-to-inosine genetic recoding is required in the adult stage nervous system for coordinated behavior in Drosophila. J. Biol. Chem. 284, 31391–31400.

32. Ryvkin, J., Bentzur, A., Shmueli, A., Tannenbaum, M., Shallom, O., Dokarker, S., Benichou, J.I.C., Levi, M., and Shohat-Ophir, G. (2021). Transcriptome Analysis of NPFR Neurons Reveals a Connection Between Proteome Diversity and Social Behavior. Front. Behav. Neurosci. 15, 628662.

33. Ingleby, L., Maloney, R., Jepson, J., Horn, R., and Reenan, R. (2009). Regulated RNA Editing and Functional Epistasis in Shaker Potassium Channels. Journal of General Physiology 133, 17–27. 10.1085/jgp.200810133.

34. Jones, A.K., Buckingham, S.D., Papadaki, M., Yokota, M., Sattelle, B.M., Matsuda, K., and Sattelle, D.B. (2009). Splice-variant- and stage-specific RNA editing of the Drosophila GABA receptor modulates agonist potency. J. Neurosci. 29, 4287–4292.

35. Olson, R.O., Liu, Z., Nomura, Y., Song, W., and Dong, K. (2008). Molecular and functional characterization of voltage-gated sodium channel variants from Drosophila melanogaster. Insect Biochemistry and Molecular Biology 38, 604–610. 10.1016/j.ibmb.2008.01.003.

36. Ryan, M.Y., Maloney, R., Fineberg, J.D., Reenan, R.A., and Horn, R. (2012). RNA editing in eag potassium channels: biophysical consequences of editing a conserved S6 residue. Channels 6, 443–452.

37. Yablonovitch, A.L., Fu, J., Li, K., Mahato, S., Kang, L., Rashkovetsky, E., Korol, A.B., Tang, H., Michalak, P., Zelhof, A.C., et al. (2017). Regulation of gene expression and RNA editing in Drosophila adapting to divergent microclimates. Nat. Commun. 8, 1570.

38. Cully, D.F., Paress, P.S., Liu, K.K., Schaeffer, J.M., and Arena, J.P. (1996). Identification of a Drosophila melanogaster glutamate-gated chloride channel sensitive to the antiparasitic agent avermectin. J. Biol. Chem. 271, 20187–20191.

39. Etter, A., Cully, D.F., Schaeffer, J.M., Liu, K.K., and Arena, J.P. (1996). An amino acid substitution in the pore region of a glutamate-gated chloride channel enables the coupling of ligand binding to channel gating. J. Biol. Chem. 271, 16035–16039.

40. Liu, W.W., and Wilson, R.I. (2013). Glutamate is an inhibitory neurotransmitter in the Drosophila olfactory system. Proc. Natl. Acad. Sci. U. S. A. 110, 10294– 10299.

41. Wolstenholme, A.J. (2012). Glutamate-gated Chloride Channels. Journal of Biological Chemistry 287, 40232–40238. 10.1074/jbc.r112.406280.

42. Kondo, S., Takahashi, T., Yamagata, N., Imanishi, Y., Katow, H., Hiramatsu, S., Lynn, K., Abe, A., Kumaraswamy, A., and Tanimoto, H. (2020). Neurochemical Organization of the Drosophila Brain Visualized by Endogenously Tagged Neurotransmitter Receptors. Cell Rep. 30, 284–297.e5.

43. Li, X., Chien, C., Han, Y., Sun, Z., Chen, X., and Dickman, D. (2021). Autocrine inhibition by a glutamate-gated chloride channel mediates presynaptic homeostatic depression. Sci Adv 7, eabj1215.

44. Groschner, L.N., Malis, J.G., Zuidinga, B., and Borst, A. (2022). A biophysical account of multiplication by a single neuron. Nature 603, 119–123.

45. Li, Y., Chen, P.-J., Lin, T.-Y., Ting, C.-Y., Muthuirulan, P., Pursley, R., Ilić, M., Pirih, P., Drews, M.S., Menon, K.P., et al. (2021). Neural mechanism of spatio-chromatic opponency in the Drosophila amacrine neurons. Curr. Biol. 31, 3040–3052.e9.

46. Zhang, R., Deng, P., Jacobson, D., and Li, J.B. (2017). Evolutionary analysis reveals regulatory and functional landscape of coding and non-coding RNA editing. PLoS Genet. 13, e1006563.

47. Sapiro, A.L., Shmueli, A., Henry, G.L., Li, Q., Shalit, T., Yaron, O., Paas, Y., Billy Li, J., and Shohat-Ophir, G. (2019). Illuminating spatial A-to-I RNA editing signatures within the brain. Proc. Natl. Acad. Sci. U. S. A. 116, 2318–2327.

48. Teufel, F., Almagro Armenteros, J.J., Johansen, A.R., Gíslason, M.H., Pihl, S.I., Tsirigos, K.D., Winther, O., Brunak, S., von Heijne, G., and Nielsen, H. (2022). SignalP 6.0 predicts all five types of signal peptides using protein language models. Nat. Biotechnol. 40, 1023–1025.

49. Jones, D.T. (1999). Protein secondary structure prediction based on position-specific scoring matrices. J. Mol. Biol. 292, 195–202.

50. Nugent, T., and Jones, D.T. (2009). Transmembrane protein topology prediction using support vector machines. BMC Bioinformatics 10, 159.

51. Buchan, D.W.A., and Jones, D.T. (2019). The PSIPRED Protein Analysis Workbench: 20 years on. Nucleic Acids Res. 47, W402–W407.

52. Drozdetskiy, A., Cole, C., Procter, J., and Barton, G.J. (2015). JPred4: a protein secondary structure prediction server. Nucleic Acids Res. 43, W389– W394.

53. Sievers, F., and Higgins, D.G. (2018). Clustal Omega for making accurate alignments of many protein sequences. Protein Sci. 27, 135–145.

54. Buck, L., and Axel, R. (1991). A novel multigene family may encode odorant receptors: a molecular basis for odor recognition. Cell 65, 175–187.

55. Vosshall, L.B., Amrein, H., Morozov, P.S., Rzhetsky, A., and Axel, R. (1999). A spatial map of olfactory receptor expression in the Drosophila antenna. Cell 96, 725–736.

56. Hallem, E.A., Ho, M.G., and Carlson, J.R. (2004). The molecular basis of odor coding in the Drosophila antenna. Cell 117, 965–979.

57. Task, D., Lin, C.-C., Vulpe, A., Afify, A., Ballou, S., Brbic, M., Schlegel, P., Raji, J., Jefferis, G., Li, H., et al. (2022). Chemoreceptor co-expression in olfactory neurons. Elife 11. 10.7554/eLife.72599.

58. Gao, Q., Yuan, B., and Chess, A. (2000). Convergent projections of Drosophila olfactory neurons to specific glomeruli in the antennal lobe. Nat. Neurosci. 3, 780– 785.

59. Mombaerts, P., Wang, F., Dulac, C., Chao, S.K., Nemes, A., Mendelsohn, M., Edmondson, J., and Axel, R. (1996). Visualizing an olfactory sensory map. Cell 87, 675–686.

60. Vosshall, L.B., Wong, A.M., and Axel, R. (2000). An olfactory sensory map in the fly brain. Cell 102, 147– 159.

61. Tanaka, N.K., Endo, K., and Ito, K. (2012). Organization of antennal lobe-associated neurons in adult Drosophila melanogaster brain. J. Comp. Neurol. 520, 4067–4130.

62. Schneider, A., Ruppert, M., Hendrich, O., Giang, T., Ogueta, M., Hampel, S., Vollbach, M., Büschges, A., and Scholz, H. (2012). Neuronal basis of innate olfactory attraction to ethanol in Drosophila. PLoS One 7, e52007.

63. Devineni, A.V., and Heberlein, U. (2009). Preferential ethanol consumption in Drosophila models features of addiction. Curr. Biol. 19, 2126–2132.

64. Wang, S.-P., Guo, W.-Y., Muhammad, S.A., Chen, R.- R., Mu, L.-L., and Li, G.-Q. (2014). Mating experience and food deprivation modulate odor preference and dispersal in Drosophila melanogaster males. J. Insect Sci. 14, 131.

65. Azanchi, R., Kaun, K.R., and Heberlein, U. (2013). Competing dopamine neurons drive oviposition choice for ethanol in Drosophila. Proc. Natl. Acad. Sci. U. S. A. 110, 21153–21158.

66. Parnas, M., Lin, A.C., Huetteroth, W., and Miesenböck, G. (2013). Odor discrimination in Drosophila: from neural population codes to behavior. Neuron 79, 932–944.

67. Fernández, M. de la P., Chan, Y.-B., Yew, J.Y., Billeter, J.-C., Dreisewerd, K., Levine, J.D., and Kravitz, E.A. (2010). Pheromonal and behavioral cues trigger male-to-female aggression in Drosophila. PLoS Biol. 8, e1000541.

68. Bentzur, A., Shmueli, A., Omesi, L., Ryvkin, J., Knapp, J.-M., Parnas, M., Davis, F.P., and Shohat-Ophir, G. (2018). Odorant binding protein 69a connects social interaction to modulation of social responsiveness in Drosophila. PLoS Genet. 14, e1007328.

69. Smith, D.P. (2007). Odor and pheromone detection in Drosophila melanogaster. Pflügers Archiv - European Journal of Physiology 454, 749–758. 10.1007/s00424-006-0190-2.

70. Wang, L., and Anderson, D.J. (2010). Identification of an aggression-promoting pheromone and its receptor neurons in Drosophila. Nature 463, 227–231.

71. Joseph, R.M., Devineni, A.V., King, I.F.G., and Heberlein, U. (2009). Oviposition preference for and positional avoidance of acetic acid provide a model for competing behavioral drives in *Drosophila*. Proceedings of the National Academy of Sciences 106, 11352–11357. 10.1073/pnas.0901419106.

72. Riffell, J.A. (2013). Neuroethology: lemon-fresh scent makes flies lay eggs. Curr. Biol. 23, R1108–R1110.

73. Lai, S.-L., Awasaki, T., Ito, K., and Lee, T. (2008). Clonal analysis of Drosophila antennal lobe neurons: diverse neuronal architectures in the lateral neuroblast lineage. Development 135, 2883–2893.

74. Rosenthal, J.J.C. (2015). The emerging role of RNA editing in plasticity. J. Exp. Biol. 218, 1812–1821.

75. Galarza-Muñoz, G., Soto-Morales, S.I., Holmgren, M., and Rosenthal, J.J.C. (2011). Physiological adaptation of an Antarctic Na+/K+-ATPase to the cold. J. Exp. Biol. 214, 2164–2174.

76. Rosenthal, J.J.C., and Seeburg, P.H. (2012). A-to-I RNA editing: effects on proteins key to neural excitability. Neuron 74, 432–439.

77. Alon, S., Garrett, S.C., Levanon, E.Y., Olson, S., Graveley, B.R., Rosenthal, J.J.C., and Eisenberg, E. (2015). The majority of transcripts in the squid nervous system are extensively recoded by A-to-I RNA editing. Elife 4. 10.7554/eLife.05198.

78. Badel, L., Ohta, K., Tsuchimoto, Y., and Kazama, H. (2016). Decoding of Context-Dependent Olfactory Behavior in Drosophila. Neuron 91, 155–167.

79. Kondoh, Y., Kaneshiro, K.Y., Kimura, K.-I., and Yamamoto, D. (2003). Evolution of sexual dimorphism in the olfactory brain of Hawaiian Drosophila. Proc. Biol. Sci. 270, 1005–1013.

80. Stockinger, P., Kvitsiani, D., Rotkopf, S., Tirián, L., and Dickson, B.J. (2005). Neural circuitry that governs Drosophila male courtship behavior. Cell 121, 795– 807.

81. Sakurai, A., Koganezawa, M., Yasunaga, K.-I., Emoto, K., and Yamamoto, D. (2013). Select interneuron clusters determine female sexual receptivity in Drosophila. Nat. Commun. 4, 1825.

82. Schlief, M.L., and Wilson, R.I. (2007). Olfactory processing and behavior downstream from highly selective receptor neurons. Nat. Neurosci. 10, 623– 630.

83. Olsen, S.R., Bhandawat, V., and Wilson, R.I. (2007). Excitatory interactions between olfactory processing channels in the Drosophila antennal lobe. Neuron 54, 89–103.

84. Brusa, R., Zimmermann, F., Koh, D.S., Feldmeyer, D., Gass, P., Seeburg, P.H., and Sprengel, R. (1995). Early-onset epilepsy and postnatal lethality associated with an editing-deficient GluR-B allele in mice. Science 270, 1677–1680.

85. Gonzalez, C., Lopez-Rodriguez, A., Srikumar, D., Rosenthal, J.J.C., and Holmgren, M. (2011). Editing of human KV1.1 channel mRNAs disrupts binding of the N-terminus tip at the intracellular cavity. Nature Communications 2. 10.1038/ncomms1446.

86. Hibbs, R.E., and Gouaux, E. (2011). Principles of activation and permeation in an anion-selective Cys-loop receptor. Nature 474, 54–60.

87. Yu, J., Zhu, H., Lape, R., Greiner, T., Du, J., Lü, W., Sivilotti, L., and Gouaux, E. (2021). Mechanism of gating and partial agonist action in the glycine receptor. Cell 184, 957–968.e21.

88. Ramaswami, G., Deng, P., Zhang, R., Anna Carbone, M., Mackay, T.F.C., and Billy Li, J. (2015). Genetic mapping uncovers cis-regulatory landscape of RNA editing. Nat. Commun. 6, 8194.

89. Kabra, M., Robie, A.A., Rivera-Alba, M., Branson, S., and Branson, K. (2013). JAABA: interactive machine learning for automatic annotation of animal behavior. Nat. Methods 10, 64–67.

90. Bentzur, A., Ben-Shaanan, S., Benichou, J.I.C., Costi, E., Levi, M., Ilany, A., and Shohat-Ophir, G. (2021). Early Life Experience Shapes Male Behavior and Social Networks in Drosophila. Curr. Biol. 31, 670.

